# scPhoenix: A Contrastive Learning-based Framework with Aux–Core Feature Disentanglement Enhances Sparse Cellular Multi-omics Translation

**DOI:** 10.1101/2025.09.27.678942

**Authors:** Chaoyu Yan, Zijian He, Lihang Ye, Shikang Zheng, Nan Lin, Guanbin Li, Weizhong Li

## Abstract

Recent advances in single-cell multi-omics co-assays and spatiotemporal sequencing technologies have provided unprecedented opportunities for systematically characterizing cellular heterogeneity. However, severe sparsity and pronounced spatial heterogeneity—hallmark features of complex diseases and tumor microenvironment—remain major obstacles in deciphering cellular multi-omics data. Here, we present scPhoenix, a contrastive learning–based framework for single-cell cross-modality translation. scPhoenix adopts a two-stage Aux–Core strategy to disentangle modality-specific feature extraction from cross-modality feature interaction. Across diverse datasets, it preserves cellular heterogeneity during translation and demonstrates significant advantages for data with high sparsity. In addition, the framework integrates a contrastive learning framework with five effective data augmentation methods tailored to single-cell data. Moreover, scPhoenix’s design supports extensions to unpaired data training and spatial multi-omics translation, enabling robust performance in scenarios with high spatial heterogeneity. scPhoenix is freely available at https://github.com/liwz-lab/scPhoenix.

## Introduction

Advances in single-cell sequencing technologies have led to the emergence of diverse single-cell multi-omics profiling methods, offering unprecedented opportunities to uncover previously unrecognized cellular heterogeneity. The subsequent breakthroughs in spatial multi-omics have further enabled the simultaneous interrogation of multiple molecular layers within cells in their native spatial context, providing a promising direction for studying tissue organization and elucidating underlying biological mechanisms. Integrative analysis for multi-omics, such as genomics, transcriptomics, epigenomics, and proteomics^1,2^, offers a more comprehensive characterization of cellular states^3-10^, enhancing our understanding of complex diseases at an unprecedented depth^2,11-14^. Although a variety of single-cell and spatial multi-omics co-assays have been developed^15-20^, they generally suffer from limitations such as low sequencing depth, high sparsity, substantial noise, and high experimental costs, compared to single-modality single-cell sequencing or bulk sequencing^21-24^. These constraints have led to an urgent need for computational approaches that can either generate more accurate paired-modality data from co-assays or translate one single modality into its paired modality.

In the field of single-cell modality translation, several methods have been proposed and progressively refined. The earliest representative, BABEL^21^, is based on an autoencoder architecture without an explicit translator. It assumes that inputs from different modalities can be mapped into a shared latent space after being processed by modality-specific encoders. Subsequently, Polarbear replaces the autoencoder with a variational autoencoder (VAE) and introduced a translator module in latent space, enabling the encoder to capture modality-specific features^25^. JAMIE adopts a dual-VAE architecture and incorporates cross-modality correspondence into the reconstruction loss to align latent spaces of different modalities^26^. UnitedNet, built on an autoencoder framework, jointly performs group identification and modality prediction tasks^27^. sciPENN employs a recurrent neural network (RNN) to translate between surface protein and RNA profiles^24^. More recently, scButterfly utilizes a dual-VAE architecture with discriminative training of the translator module for multi-omics translation, achieving state-of-the-art performance^28^.

However, current single-cell cross-modality translation approaches encounter several technical bottlenecks. First, at the data level, a major challenge lies in enabling models to learn authentic cell-type representations from datasets affected by batch effects, extreme sparsity, and low signal-to-noise ratios, while simultaneously achieving denoising and imputing missing information^28^. Second, at the framework generalization level, it remains nontrivial to extend single-cell multi-omics translation methods to spatial multi-omics contexts, such that models can adapt to scenarios with high spatial heterogeneity^29^. From a biological perspective, both severe data sparsity and pronounced spatial heterogeneity are hallmark characteristics of complex diseases^30-33^. Unfortunately, existing methods have yet to overcome these limitations.

To address these challenges, we developed scPhoenix, an unsupervised generative framework for cross-modality translation of single-cell data (Fig. 1). The model design follows the principle that modality-specific feature extraction should be handled independently due to substantial differences across modalities, whereas feature interaction requires generalization and should be addressed within a shared deep semantic space. Accordingly, the framework adopts a two-stage Aux–Core architecture that separates modality-specific feature compression from cross-modality feature interaction. In the Aux module, a masked attention network adaptively emphasizes informative features, while in the Core module, multi-head self-attention^34^ (MHSA) and multi-head cross-attention^34^ (MHCA) facilitate feature exchange between modalities at multiple semantic levels. In addition, we introduce a data augmentation framework tailored for single-cell contrastive learning^35^, along with five augmentation methods designed to enhance representation learning in single-cell data.

**Fig. 1.**
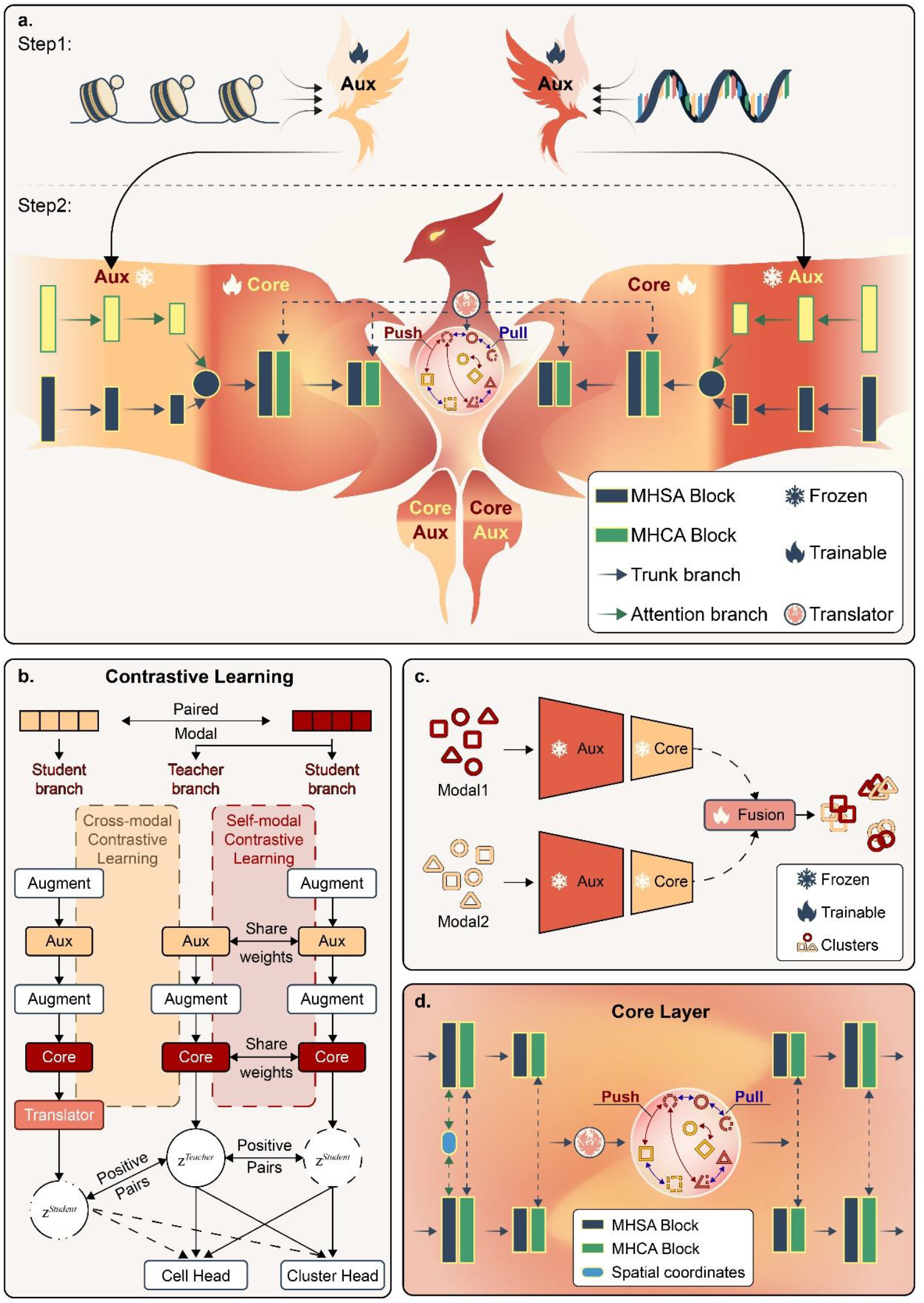
The overview architecture of scPhoenix framework. We take the translation between transcriptome and chromatin profiles as an example for illustration. **(a)** Architecture of the scPhoenix model. Paired dual-modal inputs are processed by auxiliary (Aux) and Core networks with a translator module. Training is performed in two stages: Stage 1 optimizes the Aux network with branched attention for intra-modal feature purification; Stage 2 freezes Aux weights and trains the Core (multi-head self-/cross-attention) and translator for hierarchical feature interaction and cross-modal translation. Contrastive learning aligns translated cells with their same-type counterparts while separating different cell types. **(b)** Architecture for contrastive learning. Teacher–student branches are generated by intra-modal augmentation, and translated embeddings from the paired modality serve as additional positives. **(c)** Architecture of scPhoenix-C (Cluster). This clustering variant integrates Aux/Core encoders with a lightweight VAE-based fusion module to handle unpaired multi-modal data. **(d)** Architecture of scPhoenix-S, which extends scPhoenix by integrating spatial features into the Core network for spatial multi-omics analysis.

Beyond its application to standard single-cell cross-modality translation, scPhoenix is designed with extensibility in mind. We introduce two specialized variants to address broader analytical scenarios. First, scPhoenix-C targets the integration of unpaired datasets, aiming to preserve cellular heterogeneity across diverse sources. Second, scPhoenix-S incorporates spatial awareness by disentangling spatial features from cell-intrinsic semantics, enabling its application to translation tasks in spatial multi-omics data. This design allows scPhoenix to adapt to datasets with different spatial resolutions and to operate effectively under conditions of substantial spatial perturbation.

## Results

### The model architecture of scPhenix

scPhoenix is a generative framework built upon dual-aligned autoencoders, as illustrated in Fig. 1a. The training procedure of scPhoenix follows a two-stage strategy that decouples feature extraction from feature fusion. In the first stage, we employ an auxiliary network (Aux) built upon a mask attention mechanism^36^, where the mask is specifically designed for modality-specific adaptive feature filtering and effective feature aggregation. In the second stage, the core network (Core) — comprising MHSA, MHCA, and a modality translator — is trained to perform cross-modality interaction and translation. As depicted in Fig. 1a, the complete architecture of scPhoenix consists of nine major modules, including four encoders, four decoders, and a translator. Notably, unlike conventional data integration approaches, our Aux-Core training strategy decouples feature extraction and feature interaction into two distinct stages. In this framework, encoders serve as the main network to project raw data into modality-specific latent spaces, while decoders play an auxiliary role: the Aux decoder compress features, and the Core decoder preserves the re-constructability of key features. This design facilitates cross-modality integration while maintaining modality-specific integrity. The translator, serving as a generator, enables the bidirectional mapping between latent spaces of different modalities (see Methods). To facilitate both intra- and inter-modality representation learning, we introduce a data augmentation scheme grounded in contrastive learning (Fig. 1b & Methods). We further propose a contrastive learning framework tailored for single-cell data and implement five effective methods under this framework (Fig. 2 & Methods).

**Fig. 2.**
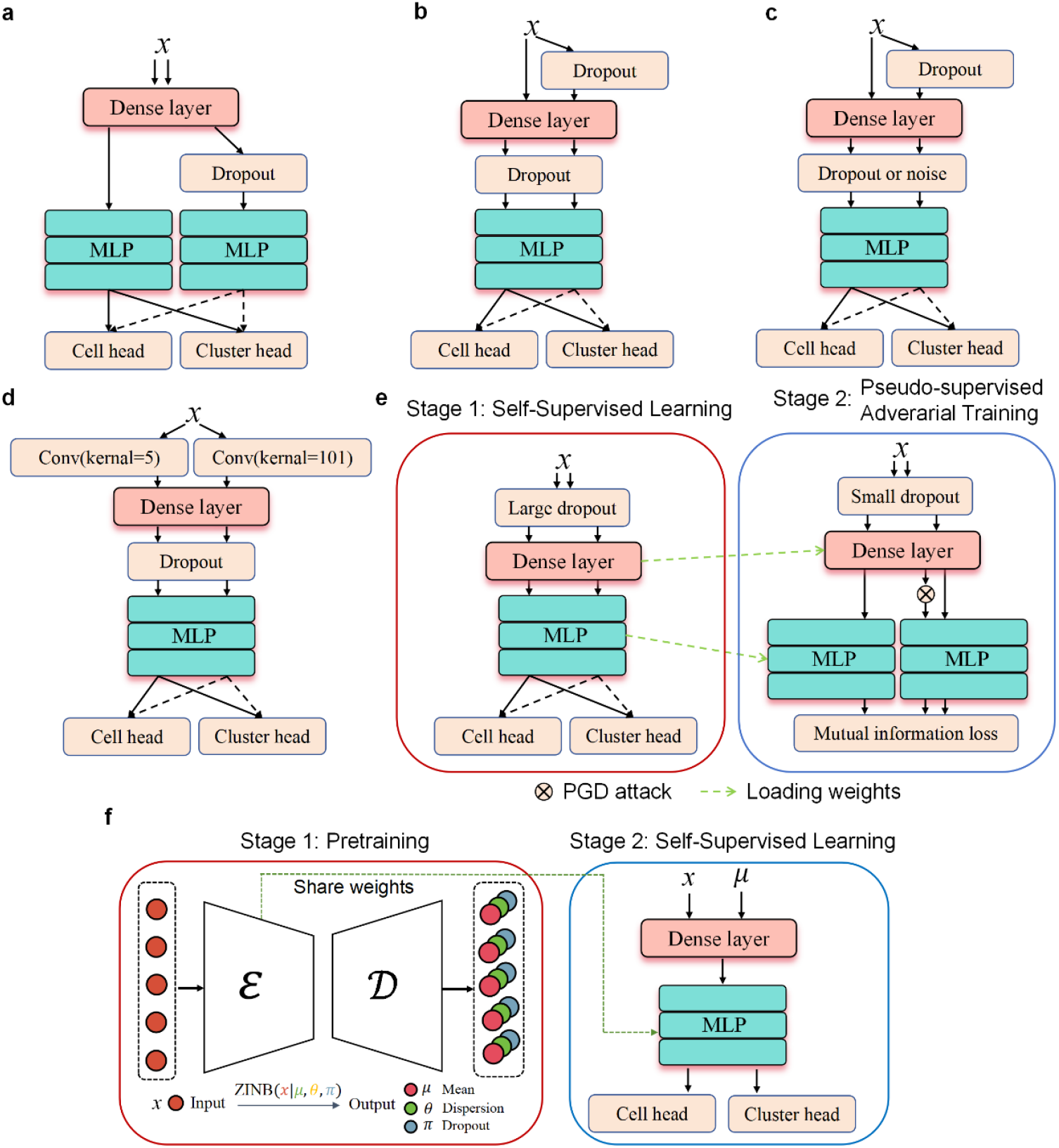
Illustration of data augmentation methods. **(a)** Baseline dual-branch model architecture. **(b)** Dropout augmentation. **(c)** Dropout and noise augmentation. **(d)** Model applying convolutional filters with varying kernel sizes to perturb input features. **(e)** Model introducing adversarial perturbations in the latent space via Projected Gradient Descent (PGD) attacks. **(f)** Model employing a Zero-Inflated Negative Binomial (ZINB) loss to denoise input data while preserving count distributions.

To enable the model to effectively handle unpaired data, we introduce a variant of scPhoenix termed scPhoenix-C (cluster), as illustrated in Fig. 1c and detailed in the Methods section. Similar to the encoder architecture in the original scPhoenix, scPhoenix-C comprises four encoders and a lightweight integration module. This design facilitates cross-modality integration and generates more accurate pseudo-pairing labels, thereby narrowing the performance gap between unpaired and paired data scenarios. Furthermore, to extend the applicability of the model to spatial multi-omics translation tasks—particularly under high spatial heterogeneity commonly observed in complex diseases—we propose another variant, scPhoenix-S (space), as shown in Fig. 1d and elaborated in the Methods section.

### The performance benchmark experiments

We employ different modality datasets to evaluate the performance of scPheonix. In the single-cell cross-modality translation task, we use scRNA-seq and scATAC-seq data from several co-assay datasets to evaluate model performance. Specifically, the CL dataset, generated using scCAT-seq from multiple cell lines, contains 549 cells across 5 cell types^18^; the MCC dataset, produced via SNARE-seq from adult mouse cerebral cortex, includes 9,190 cells annotated with 22 cell types^17^; the PBMC dataset, obtained using 10x Multiome from human peripheral blood mononuclear cells, comprises 9,631 cells spanning 19 cell types; the MDS dataset, generated with SHARE-seq from adult mouse dorsal skin, covers 32,231 cells and 22 cell types^15^; and the BMMC dataset, also profiled using 10x Multiome from human bone marrow mononuclear cells, consists of 69,249 cells covering 22 cell types^37^. Among these, the MDS dataset exhibits the highest degree of sparsity and feature dimensionality discrepancy between modalities, followed by MCC. In terms of cell-type imbalance, the BMMC dataset shows the highest level, with CL ranking second. For the single-cell clustering task, we use the MCC dataset. For the spatial multi-omics translation task, we employ datasets of varying resolution and modality types, including a human lymph node dataset generated using 10x Genomics Visium RNA and protein co-profiling technology^29^, and a mouse brain coronal co-assay dataset acquired using ATAC-RNA-seq comprising chromatin accessibility (ATAC) and transcriptomic (RNA) measurements^20^.

We conduct performance benchmark with scPheonix against existing tools and models. For the primary single-cell translation task, we compare scPhoenix against scButterfly, the current state-of-the-art (SOTA) model for single-cell translation. For the single-cell clustering task, we use MultiVI^38^ as the baseline model, while for the spatial multi-omics translation task, we employ both MultiVI and SpatialGlue^29^ as baselines. Following the evaluation protocol of scButterfly, we assess the performance of single-cell translation using four metrics: Adjusted Rand Index (ARI), Adjusted Mutual Information (AMI), Normalized Mutual Information (NMI), and Homogeneity Score (HOM), with particular emphasis on AMI, NMI, and HOM scores^39-41^. For the clustering task, we evaluate the preservation of cellular heterogeneity using AMI, NMI, HOM, and Average Silhouette Width^42^ (ASW), and assess modality integration using integration Local Inverse Simpson’s Index^42^ (iLISI), k-nearest-neighbor Batch Effect Test^42^ (kBET), and Graph Connectivity^42^ (GC). In the spatial translation task, model performance is judged by comparing spatial plots and UMAP visualizations of embedding features across different methods.

### A contrastive learning framework tailored for single-cell data

Contrastive learning has shown remarkable performance in addressing cross-modality and sparse data challenges^43,44^; however, its effectiveness in single-cell omics largely depends on appropriate data augmentation strategies, which remain a significant challenge. To address this, we propose a contrastive learning framework specifically designed for single-cell data, incorporating three core augmentation principles: (1) introducing perturbations in the latent space, (2) modifying the input receptive field, and (3) adding simulated data sampled from prior distributions. We first construct a baseline model (Fig. 2a; Methods) by integrating effective components from three representative contrastive learning studies in the single-cell domain. Building on our proposed framework, we then design five augmentation methods (Fig. 2b–f; Methods) to further enhance performance.

The baseline model employs a dual-branch architecture consisting of a teacher and a student network, both processing different views of the same data and optimized using contrastive objectives at the cell level and cluster level via corresponding projection heads. We propose five augmentation methods as follows: Method 1 applies dropout-based augmentation to the student branch and enforces shared multilayer perceptron (MLP) parameters across both branches (Fig. 2b); Method 2 further introduces Gaussian noise into the hidden layers to enhance feature perturbation (Fig. 2c); Method 3 alters the receptive fields by applying different convolutional kernels to each branch (Fig. 2d); Method 4, termed as Decoupled Adversarial Contrastive (DAC) learning^45^, leverages Projected Gradient Descent^46^ (PGD) attacks to introduce adversarial perturbations in the parameter update direction (Fig. 2e); and Method 5 simulates synthetic data following a Zero-Inflated Negative Binomial (ZINB) distribution, inspired by the denoising strategy in DCA^47^, to serve as positive pairs (Fig. 2f).

We conduct an ablation study on the MCC dataset to evaluate the effectiveness of the baseline and the proposed augmentation strategies for scRNA-seq. As shown in Table 1, our methods outperform the baseline across four clustering metrics. Specifically, Method 1 achieves the highest ARI, Method 2 yields the best AMI and NMI, and Method 4 performs best in terms of HOM, demonstrating the effectiveness of our contrastive learning framework and its tailored augmentation techniques for single-cell data analysis.

**Table 1.** Benchmarking table of data augmentation. Stratified sampling by cell type was performed to create the training set (64%), validation set (16%), and test set (20%). All augmentation methods are evaluated on 9 different splits.

| Methods | ARI | AMI | NMI | HOM |
| --- | --- | --- | --- | --- |
| **Baseline** | 0.265($\pm 0.03$) | 0.429($\pm 0.03$) | 0.449($\pm 0.03$) | 0.467($\pm 0.02$) |
| **Method 1**<br>(Dropout) | **0.473**($\pm 0.02$) | 0.600($\pm 0.03$) | 0.611($\pm 0.03$) | 0.592($\pm 0.02$) |
| **Method 2**<br>(Dropout and noise) | 0.469($\pm 0.03$) | **0.607**($\pm 0.03$) | **0.618**($\pm 0.03$) | 0.606($\pm 0.02$) |
| **Method 3**<br>(Convolution) | 0.348($\pm 0.02$) | 0.560($\pm 0.02$) | 0.575($\pm 0.02$) | 0.599($\pm 0.02$) |
| **Method 4**<br>(DAC) | 0.372($\pm 0.04$) | 0.593($\pm 0.03$) | 0.610($\pm 0.03$) | **0.653**($\pm 0.03$) |
| **Method 5**<br>(Simulation) | 0.300($\pm 0.02$) | 0.489($\pm 0.03$) | 0.506($\pm 0.03$) | 0.524($\pm 0.02$) |

DAC: Decoupled Adversarial Contrastive.

### Single-cell cross-modality translation between transcriptome and epigenome

We first compared the overall translation performance of scPhoenix and the state-of-the-art (SOTA) model scButterfly across five co-assay datasets: MCC, PBMC, CL, MDS, and BMMC. The evaluation included both latent-level translation accuracy (measured on features after the translator module) and reconstruction-based translation accuracy (measured after decoding the translated features), as shown in Fig. 3a. The overall performance score was computed as the average of AMI, NMI, and HOM. As illustrated in Fig. 3a, scPhoenix consistently outperforms scButterfly on all the datasets, with particularly larger margins on MCC, MDS, and BMMC—datasets characterized by larger sample sizes, higher sparsity, greater individual heterogeneity, and stronger batch effects^28^. These results highlight the robustness of scPhoenix, as it consistently maintains superior performance across datasets that vary widely in species, sequencing technologies, sample sizes, feature dimensionalities, batch effects, cell-type compositions, and sparsity levels.

**Fig. 3.**
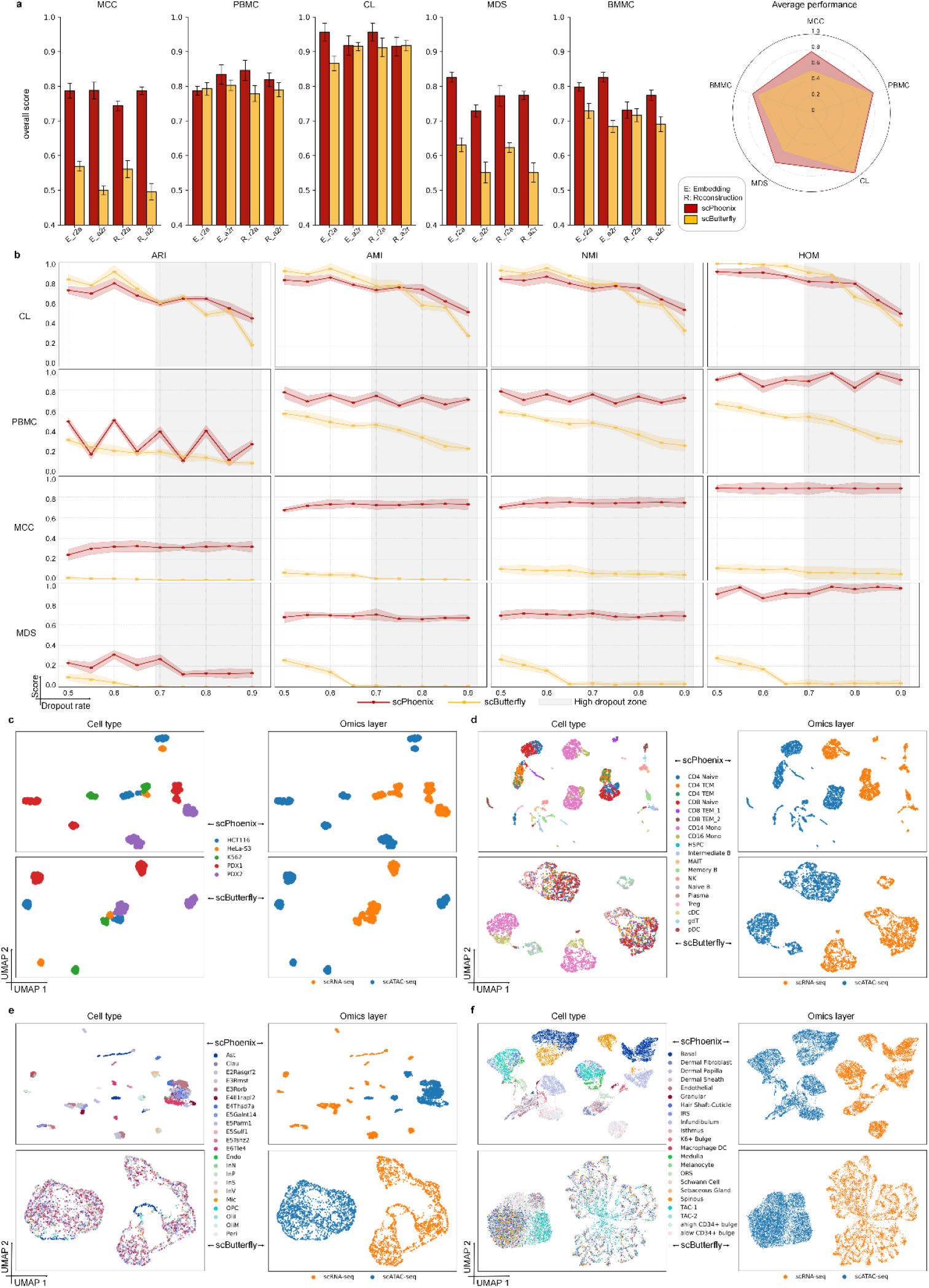
Cross-modality translation performance between the epigenome and transcriptome. **(a)** Comparison of overall translation performance across five datasets, where the overall score combines metrics from both the translated latent space and the reconstructed original space. R2A refers to translating RNA to ATAC, while A2R refers to translating ATAC to RNA. Each model was evaluated over n = 9 runs with different random seeds for data splitting. Error bars represent the mean ± standard deviation. **(b)** Performance comparison under varying dropout rates applied to the input features after preprocessing across four datasets. The red curve represents scPhoenix, and the yellow curve represents scButterfly. The shaded gray region indicates the high-dropout zone. Results are averaged over n = 9 runs with different random seeds; error bars denote mean ± standard deviation. **(c–f)** UMAP visualizations of the translated profiles under a dropout rate of 0.6, shown for four datasets: (**c**) CL, (**d**) PBMC, (**e**) MCC, and (**f**) MDS.

We further evaluated the translation performance of the model under varying levels of data sparsity on the CL, PBMC, MCC, and MDS datasets (Fig. 3b). Across all the datasets, scPhoenix consistently preserved cellular heterogeneity more effectively than scButterfly in regions with high sparsity. This performance is particularly notable in datasets such as MCC and MDS, which are characterized by high lineage diversity, continuous differentiation trajectories, complex proliferative states, and multi-dimensional heterogeneity—posing significant challenges for both feature extraction and cross-modality translation. Under highly sparse conditions, these datasets demand strong capabilities in denoising and interpolation from the model. Compared to scButterfly, scPhoenix demonstrates a clear and growing advantage as data sparsity increases and cellular heterogeneity intensifies, underscoring its robustness in handling complex, noisy, and biologically diverse single-cell data.

To further assess cellular heterogeneity preservation, we conducted visualization of the translated data under a fixed dropout rate of 0.6 across four datasets (CL, PBMC, MCC, and MDS) (Fig. 3c–f). Across all datasets, scPhoenix consistently maintained clear cell-type separation, whereas the performance of scButterfly deteriorated as dataset complexity increased. For example, in the PBMC dataset, scPhoenix exhibited partial overlap between CD4 TCM, CD4 TEM, and a subset of CD8 TEM_1 cells, but successfully distinguished the remaining 16 cell types. In contrast, scButterfly was only able to distinguish CD14 Mono, CD16 Mono, and CD8 TEM_2. Similarly, in the MCC dataset, scButterfly could only separate the Astrocyte (Ast) population, and in the MDS dataset, it distinguished the TAC-1 subtype only. In both cases, scPhoenix was able to clearly resolve the majority of cell types. These results further confirm that scPhoenix not only enables cross-modality translation but also enhances the discrimination of subtle cellular subtypes, especially in datasets with high sparsity and low inter-cell-type contrast. This capability is particularly advantageous under low sequencing depth or noisy conditions, and makes the model well-suited for analyzing complex disease microenvironments such as the tumor immune microenvironment, which often exhibit rich cellular diversity and fine-grained subtype composition.

### Single-cell unpaired data integration between transcriptome and epigenome

To enable the model to handle unpaired single-cell data, we propose a variant termed scPhoenix-C, which integrates different modalities into a shared latent space and employs Leiden clustering to generate pseudo-paired labels. This allows scPhoenix to effectively operate under unpaired modality settings. We evaluated scPhoenix-C on the MCC dataset and compared its performance to MultiVI, a representative SOTA method for unpaired integration in benchmarking. As shown in Fig. 4a, scPhoenix-C outperforms MultiVI in both cellular heterogeneity preservation and modality mixing quality, demonstrating its superior capability in learning unified representations from unpaired multimodal data. Furthermore, visualization results (Fig. 4b) show that scPhoenix-C achieves effective modality mixing, providing an intuitive confirmation of its integration performance.

**Fig. 4.**
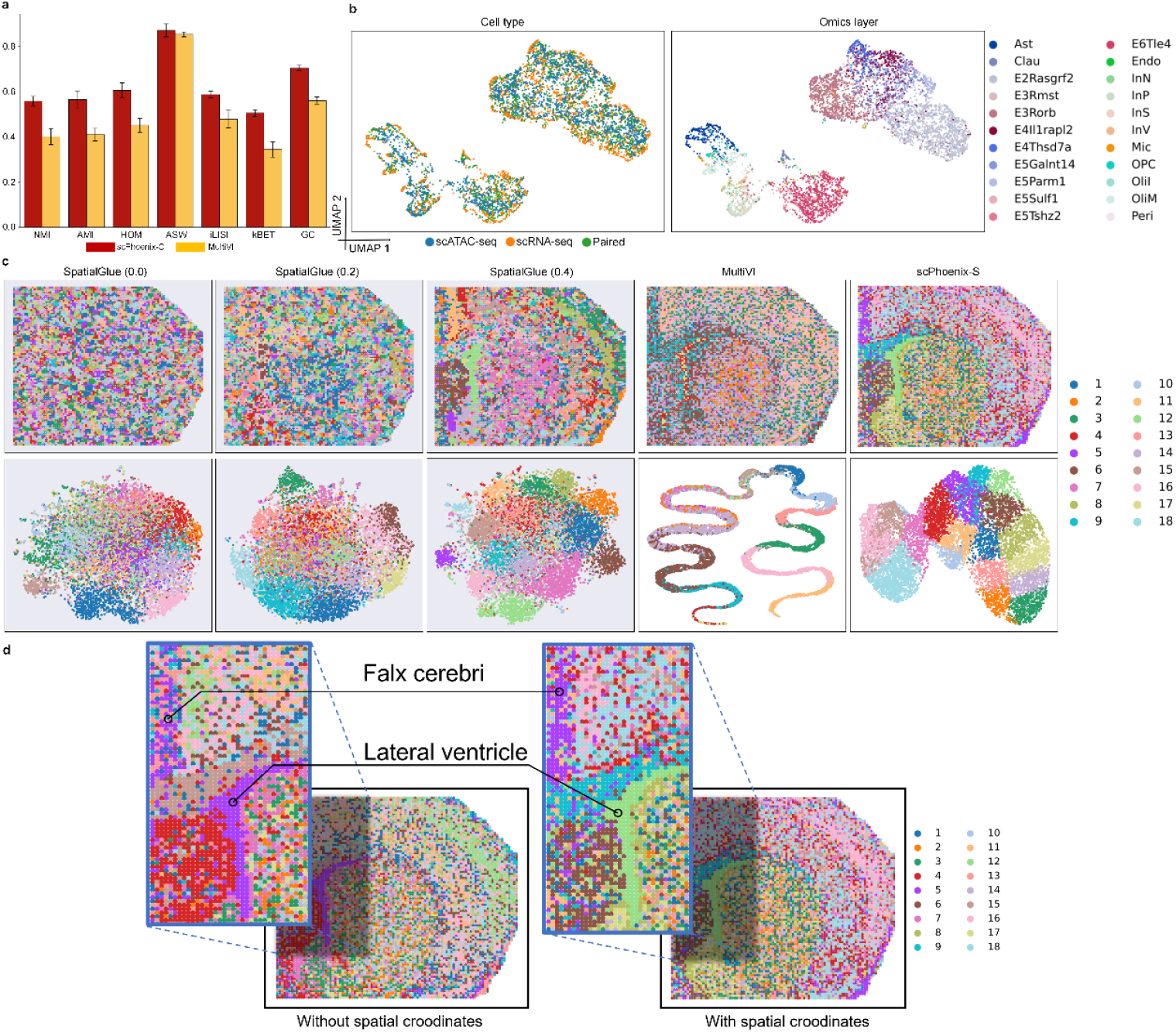
Unpaired clustering performance and spatial translation between the epigenome and transcriptome. **(a)** Bar plot comparing the performance of scPhoenix-C and MultiVI on the MCC dataset. One-third of the cells contain only scATAC-seq data, one-third only scRNA-seq data, and the remaining third are paired. Each model was evaluated over n = 9 runs with different random seeds for dataset splitting. Error bars indicate the mean ± standard deviation. The first four metrics (NMI, AMI, HOM, ASW) assess the preservation of cellular heterogeneity, while the last three metrics (iLISI, kBET, GC) evaluate cross-modality integration quality. **(b)** UMAP visualizations of the clustering results. The left panel is colored by modality, and the right panel is colored by ground truth cell types. **(c)** UMAP visualizations of embedding features (bottom) and spatial plots (top) generated by SpatialGlue, MultiVI, and scPhoenix-S. Clustering was performed using the Mclust algorithm, yielding 18 clusters. To simulate spatial disruption, the original spatial coordinates were randomly shuffled. The values in parentheses indicate the degree of spatial preservation after shuffling. **(d)** Visualization of brain functional region identification using the scPhoenix-S model, before and after the incorporation of spatial features.

### Spatial cross-modality translation

To evaluate cross-modality learning at subcellular resolution, we applied our framework to a co-assay dataset of mouse brain spatial transcriptomics and epigenomics^20^. This dataset comprises 9,215 spatial pixels, measuring 22,731 genes and 35,270 chromatin-accessible peaks, covering seven anatomical brain regions: the falx cerebri, cortex, corpus callosum, lateral ventricle, striatum, anterior commissure, and lateral septal nucleus. Within specific brain regions, cells of the same type tend to exhibit spatial aggregation; however, due to the coexistence of heterogeneous cell populations and constraints imposed by developmental and functional organization, their distribution follows a discontinuous clustering pattern. We compared the spatial mappings obtained by scPhoenix-S after latent translation with the clustering visualizations of SpatialGlue and MultiVI (Fig. 4c). To test robustness to spatial perturbation, we introduced varying levels of random noise to the original spatial coordinates. Compared with MultiVI, scPhoenix-S more clearly preserved anatomical boundaries. Notably, SpatialGlue was highly sensitive to spatial noise, and at a perturbation level of 60%, it failed to distinguish major brain regions. UMAP visualizations showed that scPhoenix-S yielded more distinct cluster boundaries than the other methods. Furthermore, we examined whether spatial information explicitly enhanced the ability of scPhoenix to delineate anatomical domains. Compared to the baseline scPhoenix, the spatial variant (scPhoenix-S) was able to distinguish between closely positioned regions such as the falx cerebri and lateral ventricle (Fig. 4d).

To assess the capability of different algorithms in capturing spatial domain-level features, we employed a human lymph node spatial transcriptomics and surface protein dataset characterized by multicellular resolution^29^. We compared the visualization performance of scPhoenix-S, scPhoenix, MultiVI, and SpatialGlue (Fig. 5). Compared with MultiVI, scPhoenix-S exhibited markedly superior delineation of cluster boundaries in the latent space as well as more accurate spatial domain localization. Furthermore, in contrast to scPhoenix, scPhoenix-S was able to identify peripheral long-range structures, such as pericapsular adipose tissue, indicating its effective utilization of spatial features (Fig. 5). Although the spatial resolution achieved by scPhoenix-S was slightly inferior to that of SpatialGlue, it nonetheless demonstrated its capacity to extract spatial domain-level features. Consistent with previous findings, SpatialGlue continued to exhibit high sensitivity to perturbations in spatial information.

**Fig. 5.**
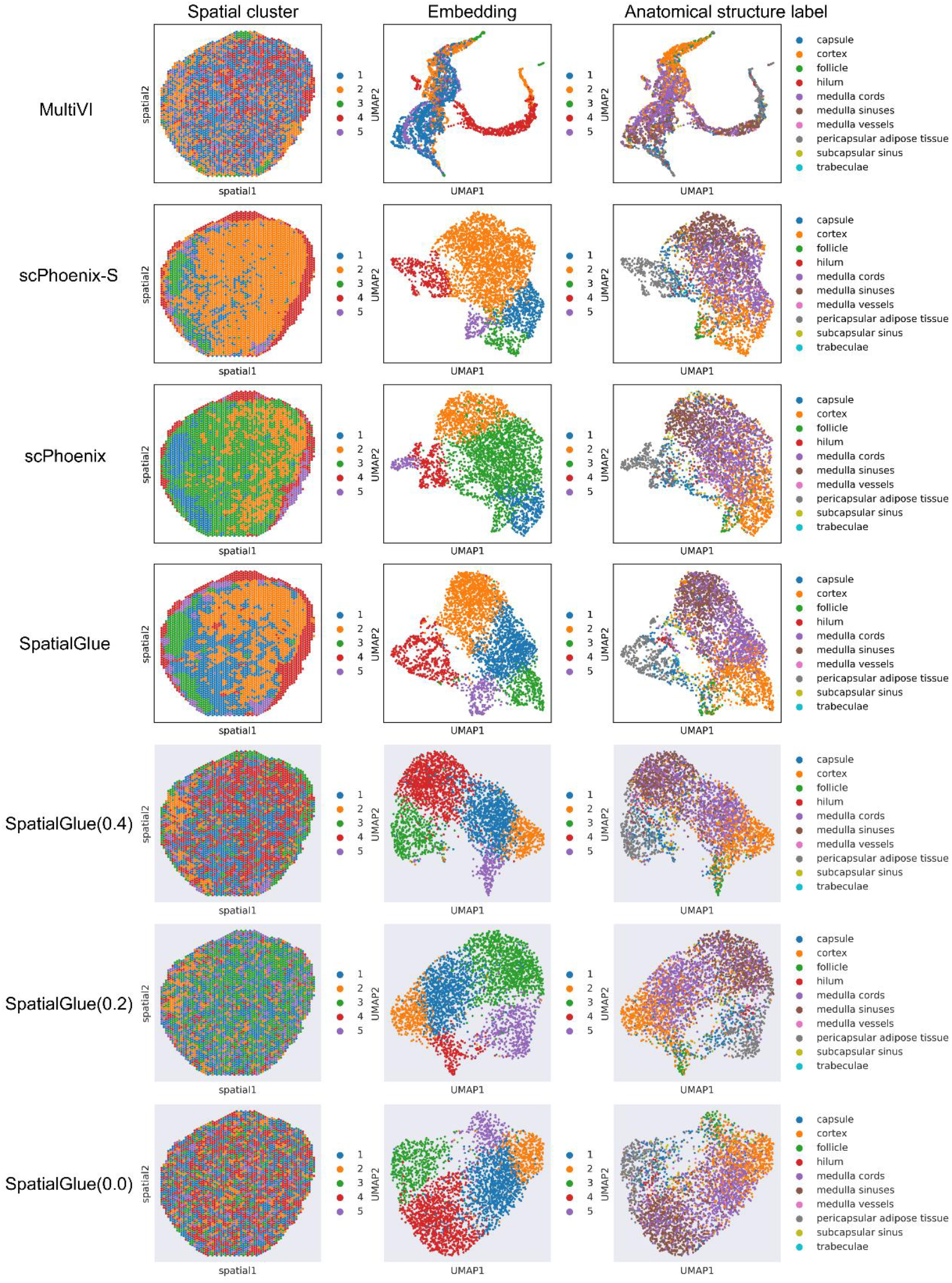
Spatial plots, UMAP visualization, and anatomical region identification on spatial transcriptomics and surface protein datasets. Latent spaces were clustered into five groups. To simulate spatial perturbation, the original spatial coordinates were randomly shuffled; values in parentheses indicate the corresponding degree of spatial preservation.

Taken together, scPhoenix-S effectively leverages spatial information to perform cross-modality translation across datasets of varying spatial resolutions. Owing to its disentangled training strategy that separates spatial features from cellular semantic representations, scPhoenix-S circumvents the common assumption in graph-based models that spatial proximity necessarily implies cellular similarity. This design makes it particularly well-suited for highly heterogeneous contexts such as the tumor immune microenvironment, where such assumptions often fail.

## Discussion

In this study, we propose scPhoenix, an unsupervised generative framework for cross-modality translation of single-cell data. Our method addresses two fundamental challenges in the single-cell and spatial multi-omics domains: (1) severe data sparsity, driven by the prevalence of dropout events and technical noise^22^, and (2) pronounced spatial heterogeneity, reflecting location-dependent variation in cellular composition and states. Biologically, these characteristics are hallmark features of complex diseases, particularly within the tumor immune microenvironment (TIME), where diverse immune cell populations interact dynamically in a spatially organized manner. From a computational standpoint, sparsity hinders the extraction of robust and informative features, whereas spatial heterogeneity increases the model’s reliance on capturing subtle, context-dependent semantic relationships between cell types, both of which present substantial obstacles to conventional analytical frameworks. However, existing single-cell translation algorithms fail to account for extreme sparsity, while most spatial omics approaches depend on the assumption that spatial proximity implies similarity— a premise that often breaks down in highly heterogeneous contexts such as TIME.

scPhoenix introduces several methodological innovations: (1) a two-stage Aux-Core architecture that disentangles feature extraction and feature fusion; (2) a contrastive learning framework tailored to the single-cell domain, along with five novel data augmentation strategies that outperform standard baselines; and (3) an MHSA+MHCA design that integrates features across different modalities and semantic depths. Extensive experiments across diverse datasets—varying in species, modality, resolution, and technical protocols—demonstrate that scPhoenix consistently outperforms existing methods in preserving cellular heterogeneity during cross-modality translation. Moreover, under simulated high-sparsity conditions, scPhoenix exhibits superior translation performance compared to baseline models.

To demonstrate the extensibility of scPhoenix beyond standard single-cell translation tasks, we developed two specialized variants tailored to broader applications. scPhoenix-C is designed for the integration of unpaired datasets, effectively preserving cellular heterogeneity across diverse sources. scPhoenix-S is a spatially aware variant that explicitly disentangles spatial features from cell-intrinsic semantics, enabling accurate translation in spatial multi-omics contexts. Comprehensive evaluations across datasets with varying spatial resolutions show that scPhoenix-S consistently outperforms existing methods in both cellular-level and domain-level translation tasks, while maintaining robustness under substantial spatial perturbations.

Nevertheless, this study has two main limitations. First, due to the lack of paired multi-omics co-assay datasets derived from tumor immune microenvironments (TIME), we only simulated the high sparsity and pronounced spatial heterogeneity characteristic of TIME rather than validating on real paired datasets. Second, the current framework has not yet been optimized for computational efficiency, which may limit its scalability for extremely large single-cell datasets.

## Methods

### Overall of scPhoenix

scPhoenix is built upon a dual-aligned autoencoder framework trained in a two-stage Aux-Core scheme. Using cross-translation between scRNA-seq and scATAC-seq as an example, the model consists of nine major components (Fig. 1a): the Aux-layer encoders for RNA and ATAC, denoted as *En*_*r*−*aux*_ and *En*_*a*−*aux*_; the Core-layer encoders for RNA and ATAC, denoted as *En*_*r*−*core*_ and *En*_*a*−*core*_; the Aux-layer decoders for RNA and ATAC, denoted as *De*_*r*−*aux*_ and *De*_*a*−*aux*_; the Core-layer decoders for RNA and ATAC, denoted as *De*_*r*−*core*_ and *De*_*a*− *core*_ ; and a translator module, denoted as *T*. Given pre-processed paired training data of scRNA-seq 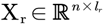, where *n* denotes the number of cells and *I*_*r*_/*I*_*a*_ denotes the feature dimensions of RNA/ATAC data, scPhoenix translates the input X_r_ to chromatin profiles by *De*_*a*−*aux*_(*De*_*a*−*core*_(*T*_*r*⟶*a*_(*En*_*r*−*core*_(*En*_*r*−*aux*_(X_r_))))), and translates the input X_a_ to transcriptome profiles by *De*_*r*−*aux*_(*De*_*r*−*core*_(*T*_*a*⟶*r*_(*En*_*a*−*core*_(*En*_*a*−*aux*_(X_a_))))). Details of each component in scPhoenix are as follows.

### Auxiliary layer network

The Aux (Auxiliary) layer constitutes the outermost component of the scPhoenix framework, functioning as a supporting module for modality-specific feature extraction and aggregation. The Aux layer adopts a mask attention architecture, where the encoder comprises a main backbone network (*E*_*main*_) and an auxiliary mask network (*E*_*branch*_) that enhances feature extraction by filtering out irrelevant features (Fig. 1a). The overall structure of the Aux layer network can thus be expressed as:

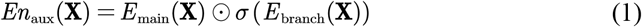

Here, *σ*(·) denotes the Sigmoid activation function, which normalizes the branch output to the range [0,1], and ⊙ denotes the Hadamard product, which applies the attention weights element-wise to the main backbone features. Both the main backbone network and the attention branch network in the Aux layer comprise three linear layers, with the final layer output dimension set to 1024. Each linear layer is followed by a Batch Normalization (BN) layer and an activation layer. Except for the final activation layer in the attention branch, which employs a Sigmoid function to normalize attention weights, all other activation layers use LeakyReLU.

### Core layer network

The Core layer network is responsible for feature interaction and cross-modality translation. Its functionality can be divided into three parts: (1) long-range interactions within the high-dimensional semantic features of each modality; (2) multi-scale cross-modality feature interactions; and (3) modal translation (Fig. 1a). The Core layer consists of five modules: *En*_*r*−*core*_, *En*_*a*−*core*_, *De*_*r*−*core*_, *De*_*a*−*core*_, and *T*. For the encoders, the input feature dimension is 1024, which is down-sampled twice to a final dimension of 256. At both the 1024-dimensional and 512-dimensional representations, a multi-head self-attention (MHSA) layer is applied to capture global and long-range feature interactions within each modality, followed by a multi-head cross-attention (MHCA) layer to learn features from the corresponding paired modality. For a given feature matrix X ∈ ℝ ^*N* × *d*^, the query, key, and value representations are computed as:

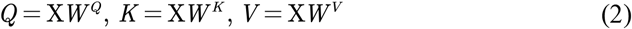

where 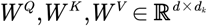 are learnable projection matrices and 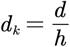 is the dimension per attention head with *h* being the number of heads. The multi-head self-attention is then formulated as:

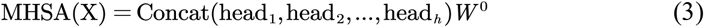

where each attention head is computed as:

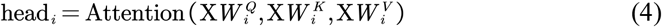

and 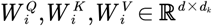 are head-specific projection matrices, 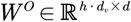 is the output projection, and *d*_*v*_ = *d*_*k*_.

For MHCA, the query is derived from the current modality 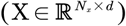, while the key and value are taken from the paired modality 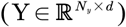:

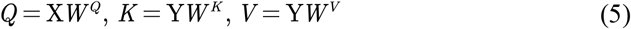

The single-head cross-attention is computed as:

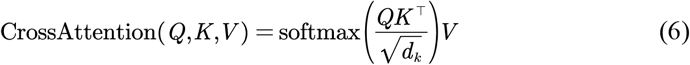

The MHCA is expressed as:

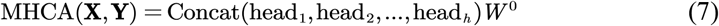

where each cross-attention head is:

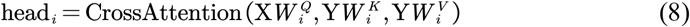

The translator module (T) consists of two fully connected layers, each with an output dimension of 256, each followed by Batch Normalization (BN) and LeakyReLU activation. The RNA representation is computed as X_*r*_ = *Linear*_2_ (*Linear*_1_(X_*a*_)).

### Contrastive learning in scPhoenix

In scPhoenix, dropout is employed as the primary data augmentation strategy (Fig. 1b). For the same modality, the model processes data through two parallel branches: a teacher branch and a student branch. The dense layer and the multilayer perceptron (MLP) correspond to the Aux and Core layers of the main network, respectively. In the projection layer design, we extend the concepts from scCobra^48^ and Contrastive Clustering^49^ by adopting a dual-head projection structure. Specifically, the original cell-level projection head (Cell Head) outputs a soft probability distribution of each cell over clusters, while the newly introduced cluster-level projection head (Cluster Head) produces a soft probability distribution of each cluster over cells.

Depending on the modality, contrastive learning can be classified into intra-modality and cross-modality paradigms. In the intra-modality setting, a positive pair consists of the latent representations of the same sample obtained from the teacher and student branches. In the cross-modality setting, a positive pair is formed by the latent representation from the teacher branch of one modality and the corresponding latent representation generated by passing the paired modality’s student branch output through a modality-specific translator.

Mathematically, we take the intra-modality training as an example. Let *N* denote the number of cells in a mini-batch, and suppose each cell undergoes one stochastic augmentation, producing 2*N* augmented samples 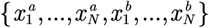. For any sample 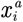, there are 2*N* − 1 total pairings: the pair 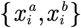 is regarded as positive, while the remaining 2*N* − 2 pairs are treated as negatives. The cell-level contrastive loss is then defined as:

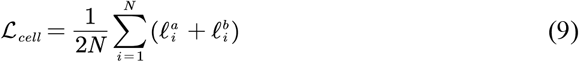

Among them,

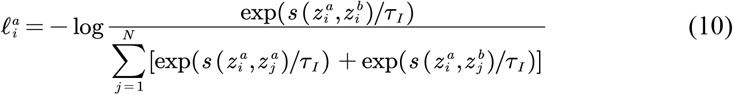

Here, *τ*_*I*_ is the temperature parameter at the cell level. Let *k*_1_,*k*_2_ ∈ {*a,b*}, and *i,j* ∈[1,*N*]. Then: 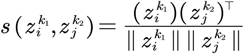, where *z*_*i*_ and *z*_*j*_ represent the hidden space features corresponding to 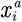 and 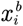 .

The cluster-level contrastive loss is:

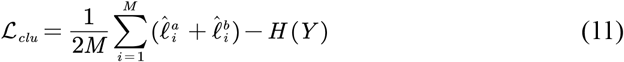

Among them,

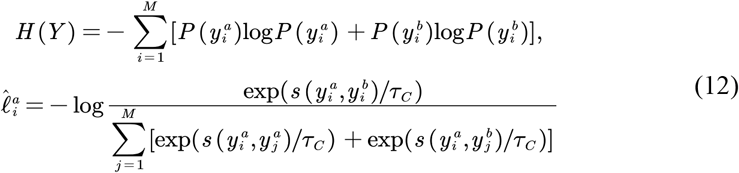

Here, *τ*_*C*_ is the temperature coefficient at the cluster level. Let *k*_1_,*k*_2_ ∈ {*a,b*} and *i,j* ∈[1,*M*]. Then: 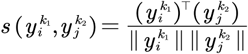 . The overall contrastive learning objective integrates both cell- and cluster-level losses with learnable weights *ω*_1_ and *ω*_2_:

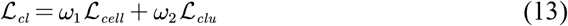

For the cross-modality setting, the loss function remains the same as in the intra-modality case, with the only difference lying in the definition of positive pairs.

### Data augmentation for contrastive learning

We conducted a unified evaluation on scRNA-seq data, assessing the clustering performance of latent features learned by the model. For all experiments, the dense layer dimension was fixed at 1024, the encoder was implemented as a three-layer MLP, and the dimensions of the cell head and cluster head were kept identical.

### Baseline model (Fig. 2a)

The input data are first passed through a dense layer and then split into two branches. One branch is processed by an MLP, while the other undergoes dropout before being fed into another MLP with the same architecture but different parameters. The resulting latent features from the two branches are then passed through the cell head and cluster head, respectively, to compute the contrastive loss for model training.

### Dropout-based data augmentation strategy (Fig. 2b)

For clarity, the branch with smaller perturbations is referred to as the teacher branch (left), whereas the branch with larger perturbations is referred to as the student branch (right). After model training, the teacher branch is the focus of this study, and the latent features obtained from the MLP layer of the teacher branch serve as the target low-dimensional representations.

### Dropout with noise-based data augmentation strategy (Fig. 2c)

After passing through the dense layer, the data are randomly subjected to either dropout or Gaussian noise with randomly assigned weights.

### Convolution-based data augmentation strategy (Fig. 2d)

In the teacher branch, the data are processed by a one-dimensional convolution with a kernel size of 5, whereas in the student branch, a one-dimensional convolution with a kernel size of 101 is applied. The core idea is that the smaller kernel in the teacher branch enables the capture of more fine-grained features from the raw data, while the larger kernel in the student branch facilitates the extraction of more global features.

### Interpretable adversarial contrastive data augmentation strategy (Fig. 2e)

The core idea is to enhance perturbations based on the parameter update direction, following a two-step scheme. A teacher model is first trained and subsequently used to guide the student model in performing contrastive learning. In the second step, the loss function incorporates both the similarity between the embeddings obtained from the teacher model and those from the student model, as well as the similarity between the teacher model and the adversarially perturbed student model generated via the PGD attack:

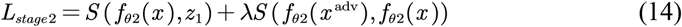

where:

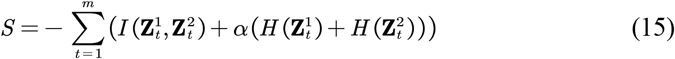

Here, *I* denotes mutual information, and *H* denotes information entropy. The inclusion of information entropy aims to prevent the model from mapping all samples to the same point, thus avoiding mode collapse.

The adversarial perturbation is generated as follows:

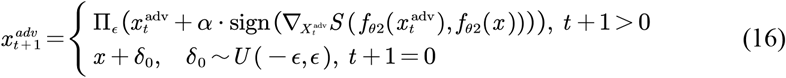

where 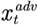 is the adversarial sample at step *t, ϵ* is the maximum allowable perturbation magnitude, and *α* is the step size of each perturbation. The operator Π _*ϵ*_ denotes the projection that constrains the perturbation within the *ϵ*− ball. *δ*_0_ represents the initial random perturbation, uniformly sampled from [− *ϵ, ϵ*].

### ZINB-based data augmentation strategy (Fig. 2f)

Following the idea of DCA, the core assumption is that the input data follow a zero-inflated negative binomial (ZINB) distribution. An autoencoder is then employed to reconstruct the outputs *μ, θ*, and *π* .

The pre-trained reconstructions *μ* and *θ*, together with the original inputs, are used as positive sample pairs for contrastive learning. The reconstruction loss is defined as follows:

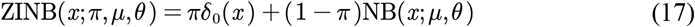

### Network architecture of scPhoenix-C

The scPhoenix-C architecture comprises three components: modality-specific Aux networks, Core networks, and a Fusion module (Fig. 1c). The Aux networks are identical to those in scPhoenix. In the Core networks, because the input modalities are unmatched, the original MHCA modules are replaced with the MHSA modules. The Fusion module is implemented as a compact variational autoencoder (VAE). Similar to MultiVI, we assume that the latent space of each modality follows a multivariate Gaussian distribution, and we employ the symmetric Jeffrey’s divergence between distributions in the evidence lower bound (ELBO) to regularize the distance between modalities in the latent space. The input to the Fusion module is 256-dimensional, with the mean layer dimension set to 16. During training, we adopt a staged strategy, sequentially training the Aux, Core, and Fusion networks.

### Network architecture of scPhoenix-S

In scPhoenix-S (Fig. 1d), spatial features are incorporated into the Core network of scPhoenix to guide cell-level semantic feature alignment across modalities in a contrastive learning framework. For spatial feature generation, the cell coordinates (*u*_*n*_, *v*_*n*_) are first normalized, after which logarithmically spaced frequencies are computed to construct the spatial embeddings:

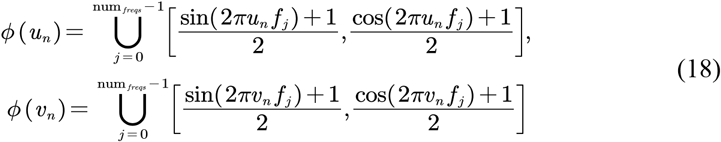

where:

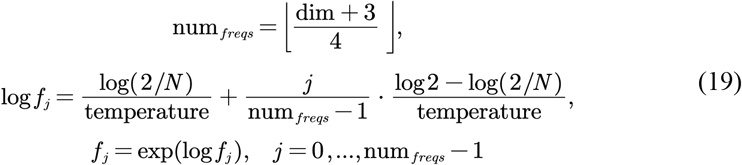

Finally, only the first K dimensions of the spatial features are retained.

For spatial feature guidance, the generated spatial features are subjected to dropout and then concatenated directly to the semantic features. In this process, the teacher branches of different modalities share the same spatial features, and likewise, the student branches of different modalities also share the same spatial features.

### Training procedure

scPhoenix adopts a two-stage Aux–Core training scheme. In the Aux stage, the encoders and decoders are optimized in a self-supervised manner: the RNA modality employs mean squared error (MSE) as the reconstruction loss, whereas the ATAC modality uses binary cross-entropy (BCE). Training in this stage is performed using the Adam optimizer with a learning rate of 0.0001, for up to 150 epochs, with early stopping applied if no improvement is observed for 10 consecutive epochs.

In the Core layer, the loss function is defined as:

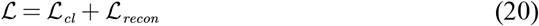

where:

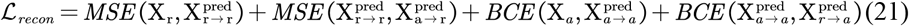

and:

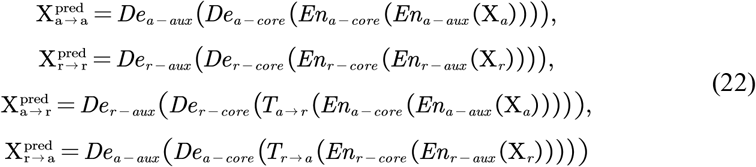

By default, scPhoenix is optimized using Adam with a learning rate of 0.0001 and minibatch size of 128. It should be noted that the optimal performance for different datasets was not necessarily achieved at the same learning rate. We evaluated five learning rates (0.001, 0.0003, 0.0001, 0.00003, and 0.00001) and observed that the model exhibited the most stable performance at 0.0001. For non-extremely sparse datasets, the initial dropout applied to the student branch was set to less than 50%, whereas for extremely sparse datasets, the initial dropout was set to less than 25%. As in scButterfly, we used 64% of the data for training, 16% for validation, and the remaining 20% for testing. In the Core layer, the model was trained for 300 epochs with an early stopping patience of 50 epochs.

### Data pre-processing

Following the scButterfly pre-processing pipeline, for the count metrics of scRNA-seq profile, we first performed normalization followed by logarithmic transformation, and selected the top 3,000 highly variable genes as the optimal feature set. For the count metrics of scATAC-seq profile, we first applied binarization, followed by peak filtering, term frequency–inverse document frequency^40,50,51^ (TF-IDF) transformation, and finally scaling. For the count metrics of proteome profiles, we performed centered log-ratio^52^ (CLR) transformation.

### Evaluation metrics

Cell heterogeneity preservation in translated profiles was evaluated through clustering analysis. Translated profiles were first reduced to 50 dimensions using principal component analysis (PCA), followed by clustering with the Leiden algorithm^53^ (resolution = 1). Clustering quality was quantified using adjusted Rand index (ARI), adjusted mutual information (AMI), normalized mutual information (NMI), and homogeneity (HOM). Given that most single-cell datasets exhibit imbalanced cell populations, AMI is generally more suitable for such cases, whereas ARI is preferred for nearly balanced clusters. The degree of cross-modality integration was evaluated using integration Local Inverse Simpson’s Index (iLISI), k-nearest-neighbor Batch Effect Test (kBET), and Graph Connectivity (GC) as metrics.

### Visualization

For data visualization, we performed PCA to reduce the dimensionality of translated profiles to 50 and then adopted the UMAP method to further reduce the dimension to two.

## Code availability

scPhoenix is open-source for the research community and can be accessible at https://github.com/liwz-lab/scPhoenix.

## Data availability

All datasets analysed in this study are publicly available for the research community. For single-cell omics data, the CL dataset was obtained from a previously published study^18^ (https://www.nature.com/articles/s41467-018-08205-7). The MCC dataset is deposited in the Gene Expression Omnibus (GEO) (GSE126074^17^, (https://www.ncbi.nlm.nih.gov/geo/query/acc.cgi?acc=GSE126074). The PBMC dataset can be accessed from the 10x Genomics website (https://support.10xgenomics.com/single-cell-gene-expression/datasets/3.0.0/pbmc_10k_v3). The MDS dataset is available in GEO (GSE140203^15^, https://www.ncbi.nlm.nih.gov/geo/query/acc.cgi?acc=GSE140203), and the BMMC dataset is also available in GEO (GSE194122^37^, https://www.ncbi.nlm.nih.gov/geo/query/acc.cgi?acc=GSE194122). For spatial multi-omics data, the 10x Visium human lymph node dataset is accessible in GEO (GSE263617, https://www.ncbi.nlm.nih.gov/geo/query/acc.cgi?acc=GSE263617). The spatial epigenome–transcriptome mouse brain dataset is available through the AtlasXplore platform^20^ (https://web.atlasxomics.com/visualization/Fan/).

## Acknowledgements

This work was supported by the grants of National Natural Science Foundation of China (92474107), National Key R&D Program of China (2021YFF1200903), Guangdong Basic and Applied Basic Research Foundation of China (2022B1515120077), and Major Project of Guangzhou National Laboratory of China (GZNL2024A01003).

## References

1 Flynn, E., Almonte-Loya, A. & Fragiadakis, G. K. Single-Cell Multiomics. Annu Rev Biomed Data Sci 6, 313–337 (2023). 10.1146/annurev-biodatasci-020422-050645

2 Baysoy, A., Bai, Z., Satija, R. & Fan, R. The technological landscape and applications of single-cell multi-omics. Nat Rev Mol Cell Biol 24, 695–713 (2023). 10.1038/s41580-023-00615-w

3 Hao, Y. et al. Integrated analysis of multimodal single-cell data. Cell 184, 3573–3587 e3529 (2021). 10.1016/j.cell.2021.04.048

4 Gayoso, A. et al. Joint probabilistic modeling of single-cell multi-omic data with totalVI. Nat Methods 18, 272–282 (2021). 10.1038/s41592-020-01050-x

5 Zhang, L., Zhang, J. & Nie, Q. DIRECT-NET: An efficient method to discover cis-regulatory elements and construct regulatory networks from single-cell multiomics data. Sci Adv 8, eabl7393 (2022). 10.1126/sciadv.abl7393

6 Kartha, V. K. et al. Functional inference of gene regulation using single-cell multi-omics. Cell Genom 2 (2022). 10.1016/j.xgen.2022.100166

7 Li, C., Virgilio, M. C., Collins, K. L. & Welch, J. D. Multi-omic single-cell velocity models epigenome-transcriptome interactions and improves cell fate prediction. Nat Biotechnol 41, 387–398 (2023). 10.1038/s41587-022-01476-y

8 Gorin, G., Svensson, V. & Pachter, L. Protein velocity and acceleration from single-cell multiomics experiments. Genome Biol 21, 39 (2020). 10.1186/s13059-020-1945-3

9 Gayoso, A. et al. Deep generative modeling of transcriptional dynamics for RNA velocity analysis in single cells. Nat Methods 21, 50–59 (2024). 10.1038/s41592-023-01994-w

10 La Manno, G. et al. RNA velocity of single cells. Nature 560, 494–498 (2018). 10.1038/s41586-018-0414-6

11 Gong, D., Arbesfeld-Qiu, J. M., Perrault, E., Bae, J. W. & Hwang, W. L. Spatial oncology: Translating contextual biology to the clinic. Cancer Cell 42, 1653–1675 (2024). 10.1016/j.ccell.2024.09.001

12 Tirosh, I. & Suva, M. L. Cancer cell states: Lessons from ten years of single-cell RNA-sequencing of human tumors. Cancer Cell 42, 1497–1506 (2024). 10.1016/j.ccell.2024.08.005

13 Papalexi, E. & Satija, R. Single-cell RNA sequencing to explore immune cell heterogeneity. Nat Rev Immunol 18, 35–45 (2018). 10.1038/nri.2017.76

14 Ravi, V. M. et al. Spatially resolved multi-omics deciphers bidirectional tumor-host interdependence in glioblastoma. Cancer Cell 40, 639–655 e613 (2022). 10.1016/j.ccell.2022.05.009

15 Ma, S. et al. Chromatin Potential Identified by Shared Single-Cell Profiling of RNA and Chromatin. Cell 183, 1103–1116 e1120 (2020). 10.1016/j.cell.2020.09.056

16 Cao, J. et al. Joint profiling of chromatin accessibility and gene expression in thousands of single cells. Science 361, 1380–1385 (2018). 10.1126/science.aau0730

17 Chen, S., Lake, B. B. & Zhang, K. High-throughput sequencing of the transcriptome and chromatin accessibility in the same cell. Nat Biotechnol 37, 1452–1457 (2019). 10.1038/s41587-019-0290-0

18 Liu, L. et al. Deconvolution of single-cell multi-omics layers reveals regulatory heterogeneity. Nat Commun 10, 470 (2019). 10.1038/s41467-018-08205-7

19 Liu, Y. et al. High-plex protein and whole transcriptome co-mapping at cellular resolution with spatial CITE-seq. Nature Biotechnology 41, 1405–1409 (2023).

20 Zhang, D. et al. Spatial epigenome-transcriptome co-profiling of mammalian tissues. Nature 616, 113–122 (2023). 10.1038/s41586-023-05795-1

21 Wu, K. E., Yost, K. E., Chang, H. Y. & Zou, J. BABEL enables cross-modality translation between multiomic profiles at single-cell resolution. Proc Natl Acad Sci U S A 118 (2021). 10.1073/pnas.2023070118

22 Argelaguet, R., Cuomo, A. S. E., Stegle, O. & Marioni, J. C. Computational principles and challenges in single-cell data integration. Nat Biotechnol 39, 1202–1215 (2021). 10.1038/s41587-021-00895-7

23 Yang, K. D. et al. Multi-domain translation between single-cell imaging and sequencing data using autoencoders. Nat Commun 12, 31 (2021). 10.1038/s41467-020-20249-2

24 Lakkis, J. et al. A multi-use deep learning method for CITE-seq and single-cell RNA-seq data integration with cell surface protein prediction and imputation. Nat Mach Intell 4, 940–952 (2022). 10.1038/s42256-022-00545-w

25 Zhang, R., Meng-Papaxanthos, L., Vert, J. P. & Noble, W. S. Multimodal Single-Cell Translation and Alignment with Semi-Supervised Learning. J Comput Biol 29, 1198–1212 (2022). 10.1089/cmb.2022.0264

26 Kalafut, N. C., Huang, X. & Wang, D. Joint variational autoencoders for multimodal imputation and embedding. Nat Mach Intell 5, 631–642 (2023). 10.1038/s42256-023-00663-z

27 Tang, X. et al. Explainable multi-task learning for multi-modality biological data analysis. Nat Commun 14, 2546 (2023). 10.1038/s41467-023-37477-x

28 Cao, Y. et al. scButterfly: a versatile single-cell cross-modality translation method via dual-aligned variational autoencoders. Nat Commun 15, 2973 (2024). 10.1038/s41467-024-47418-x

29 Long, Y. et al. Deciphering spatial domains from spatial multi-omics with SpatialGlue. Nature Methods 21, 1658–1667 (2024). 10.1038/s41592-024-02316-4

30 Yates, J. & Van Allen, E. M. New horizons at the interface of artificial intelligence and translational cancer research. Cancer Cell 43, 708–727 (2025).

31 Qian, J. et al. Identification and characterization of cell niches in tissue from spatial omics data at single-cell resolution. Nat Commun 16, 1693 (2025). 10.1038/s41467-025-57029-9

32 Bied, M., Ho, W. W., Ginhoux, F. & Bleriot, C. Roles of macrophages in tumor development: a spatiotemporal perspective. Cell Mol Immunol 20, 983–992 (2023). 10.1038/s41423-023-01061-6

33 de Visser, K. E. & Joyce, J. A. The evolving tumor microenvironment: From cancer initiation to metastatic outgrowth. Cancer Cell 41, 374–403 (2023). 10.1016/j.ccell.2023.02.016

34 Vaswani, A. et al. Attention is all you need. Advances in neural information processing systems 30 (2017).

35 Tao, L., Wang, X. & Yamasaki, T. An Improved Inter-Intra Contrastive Learning Framework on Self-Supervised Video Representation. IEEE Transactions on Circuits and Systems for Video Technology 32, 5266–5280 (2022). 10.1109/TCSVT.2022.3141051

36 He, K. M., Zhang, X. Y., Ren, S. Q. & Sun, J. Deep Residual Learning for Image Recognition. Proc Cvpr Ieee, 770–778 (2016). 10.1109/Cvpr.2016.90

37 Luecken, M. D. et al. in 35th conference on neural information processing systems (NeurIPS 2021) track on datasets and benchmarks.

38 Ashuach, T. et al. MultiVI: deep generative model for the integration of multimodal data. Nat Methods 20, 1222–1231 (2023). 10.1038/s41592-023-01909-9

39 Chen, H. et al. Assessment of computational methods for the analysis of single-cell ATAC-seq data. Genome Biol 20, 241 (2019). 10.1186/s13059-019-1854-5

40 Chen, S. et al. RA3 is a reference-guided approach for epigenetic characterization of single cells. Nat Commun 12, 2177 (2021). 10.1038/s41467-021-22495-4

41 Shengquan, C., Boheng, Z., Xiaoyang, C., Xuegong, Z. & Rui, J. stPlus: a reference-based method for the accurate enhancement of spatial transcriptomics. Bioinformatics 37, i299–i307 (2021). 10.1093/bioinformatics/btab298

42 Luecken, M. D. et al. Benchmarking atlas-level data integration in single-cell genomics. Nat Methods 19, 41–50 (2022). 10.1038/s41592-021-01336-8

43 Radford, A. et al. in International conference on machine learning. 8748-8763 (PmLR).

44 He, K. et al. in Proceedings of the IEEE/CVF conference on computer vision and pattern recognition. 16000–16009.

45 Zhang, C. et al. in European Conference on Computer Vision. 725–742 (Springer).

46 Madry, A., Makelov, A., Schmidt, L., Tsipras, D. & Vladu, A. Towards deep learning models resistant to adversarial attacks. arXiv preprint 1706.06083 (2017).

47 Eraslan, G., Simon, L. M., Mircea, M., Mueller, N. S. & Theis, F. J. Single-cell RNA-seq denoising using a deep count autoencoder. Nat Commun 10, 390 (2019). 10.1038/s41467-018-07931-2

48 Zhao, B., Song, K., Wei, D.-Q., Xiong, Y. & Ding, J. scCobra allows contrastive cell embedding learning with domain adaptation for single cell data integration and harmonization. Communications Biology 8, 233 (2025). 10.1038/s42003-025-07692-x

49 Li, Y. et al. in Proceedings of the AAAI conference on artificial intelligence. 8547–8555.

50 Chen, X. et al. Cell type annotation of single-cell chromatin accessibility data via supervised Bayesian embedding. Nature Machine Intelligence 4, 116–126 (2022).

51 Chen, S., Wang, R., Long, W. & Jiang, R. ASTER: accurately estimating the number of cell types in single-cell chromatin accessibility data. Bioinformatics 39 (2023). 10.1093/bioinformatics/btac842

52 Stuart, T. et al. Comprehensive Integration of Single-Cell Data. Cell 177, 1888–1902 e1821 (2019). 10.1016/j.cell.2019.05.031

53 Traag, V. A., Waltman, L. & van Eck, N. J. From Louvain to Leiden: guaranteeing well-connected communities. Sci Rep 9, 5233 (2019). 10.1038/s41598-019-41695-z

